# FishExp: a comprehensive database and analysis platform for gene expression and alternative splicing of fish species

**DOI:** 10.1101/2021.07.01.450804

**Authors:** Suxu Tan, Wenwen Wang, Wencai Jie, Jinding Liu

**Author notes:** Corresponding authors (J.L.).

## Abstract

The publicly archived RNA-seq data has grown exponentially, but its valuable information has not yet been fully discovered and utilized, especially for alternative splicing. This is true for fish species, which play important roles in ecology, research, and the food industry. To mitigate this, we present FishExp, a web-based data platform covering gene expression and alternative splicing in 26,081 RNA-seq experiments from 44 fishes. In addition to searching by gene identifiers and symbols, FishExp allows users to query the data using various functional terms and BLAST alignment. Notably, the user can customize experiments and tools to perform differential/specific expression and alternative splicing analysis, provided with functional enrichments. The results of retrieval and analysis can be visualized on the gene-, transcript- and splicing event-level webpage in a highly interactive and intuitive manner. The manually curated sample information, uniform data processing and visualization tools make it efficient for users to gain new insights from these large datasets. All data in FishExp can be downloaded for more in-depth analysis. FishExp is freely accessible at https://bioinfo.njau.edu.cn/fishExp.

## Introduction

Fishes are the largest group of vertebrates, with over 34,000 species [1], more than all other vertebrate species combined. They are the earliest vertebrates on Earth and have been evolved for more than 500 million years [2,3]. Living in a variety of habitats globally, fishes play vital structural and functional roles in the aquatic ecosystem. With the wide range of time and geographical scale, they exhibit extremely high levels of biodiversity in terms of morphology, behavior, ecology, and among others. Fish species have been utilized as excellent models in studies of development, physiology, behavior, toxicology, evolution, genetics, etc. For instance, zebrafish and medaka are valuable models for studying human genetics and disease [4–6]. In addition, many fish species serve as important food sources in fishery and aquaculture [7].

With the advancement of high-throughput sequencing technologies, fish research has entered the era of omics and profoundly revolutionized our understanding of biology, diversity and disease [8]. So far, transcriptomic studies mainly focused on differential gene expression, with negligence of many other crucial layers of regulations such as splicing. Alternative splicing (AS) is the mechanism by which a single pre-mRNA molecule generates different mature mRNAs (transcripts or isoforms), enhancing proteomic diversity and gene expression modulation. Studies have demonstrated its prevalence and importance in eukaryotes, particularly in vertebrate species. For instance, nearly all multi-exonic genes in human exhibit AS events [9,10]. AS functions in a cell-, tissue-, or condition-specific manner, and plays key roles in development, disease and stress response [11].

Efficient retrieval and display of such important information can improve our understanding of gene regulatory networks and facilitate genome annotation and future functional research. A number of AS databases have been developed and become increasingly useful, including TCGA SpliceSeq [12], ASpedia [13], VastDB [14], ASlive [15] and MeDAS [16]. These databases, however, typically accommodate a small number of model organisms or provide specialized scope of knowledge and analysis, and none of them is specifically designed for fish species. Due to the important role of fish and the limited utilization of existing resources, there is an urgent need to construct a comprehensive platform for a large number of fish species. We thereby created FishExp, a user-friendly, highly interactive database and analysis platform for fishes. We developed a uniform bioinformatic pipeline to integrate genomic/transcriptomic data with comprehensive metadata to systematically analyze gene expression/AS profiles of 44 fishes. Genome annotations of most fish species were greatly improved, and many novel transcripts and splicing events were discovered and displayed. Furthermore, the user is allowed to flexibly customize RNA-seq experiments/studies of interest and examine differentially expressed genes (DEG) and differentially alternatively spliced (DAS) events/genes between treatments, tissues, developmental stages, etc. During this process, functional enrichment analysis will be automatically performed according to user-defined tools and parameters. This study provides an added-value resource for public repertoire and offers convenient tools of retrieval, analysis and visualization for the fish genome research community.

## Materials and methods

### Data collection and database content

The reference genome assemblies and original annotation of fish species were collected from Ensembl, alternatively from NCBI. RNA-seq data were collected from Sequence Read Archive (SRA) database by querying the SRA metadata (as of July, 2020) [17]. Illumina sequencing data was exclusively collected due to its ubiquity and high base quality. A total of 26,081 high-quality RNA-seq experiments of 664 studies from 44 fish species were harbored by FishExp (Table 1).

**Table 1.**
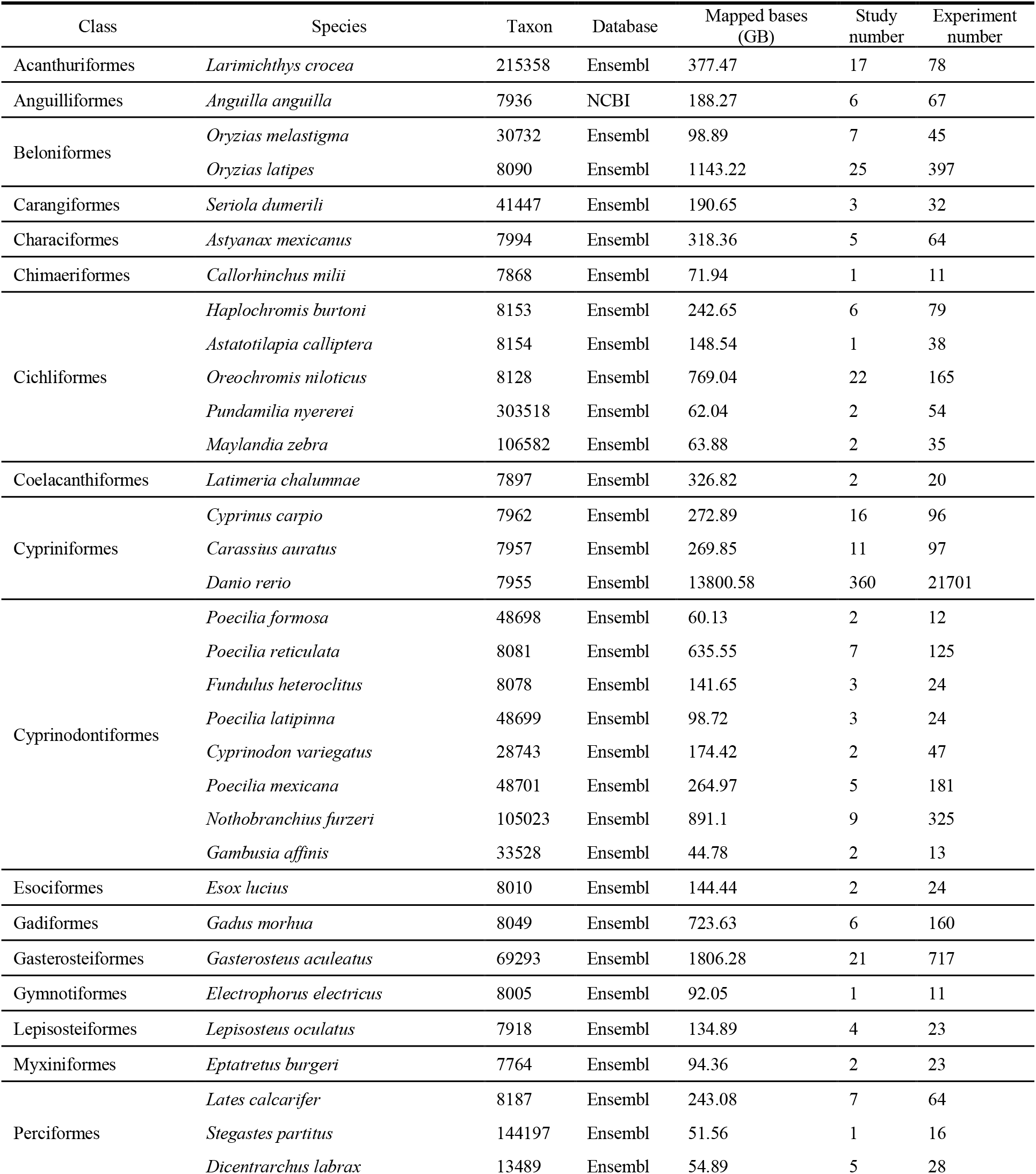
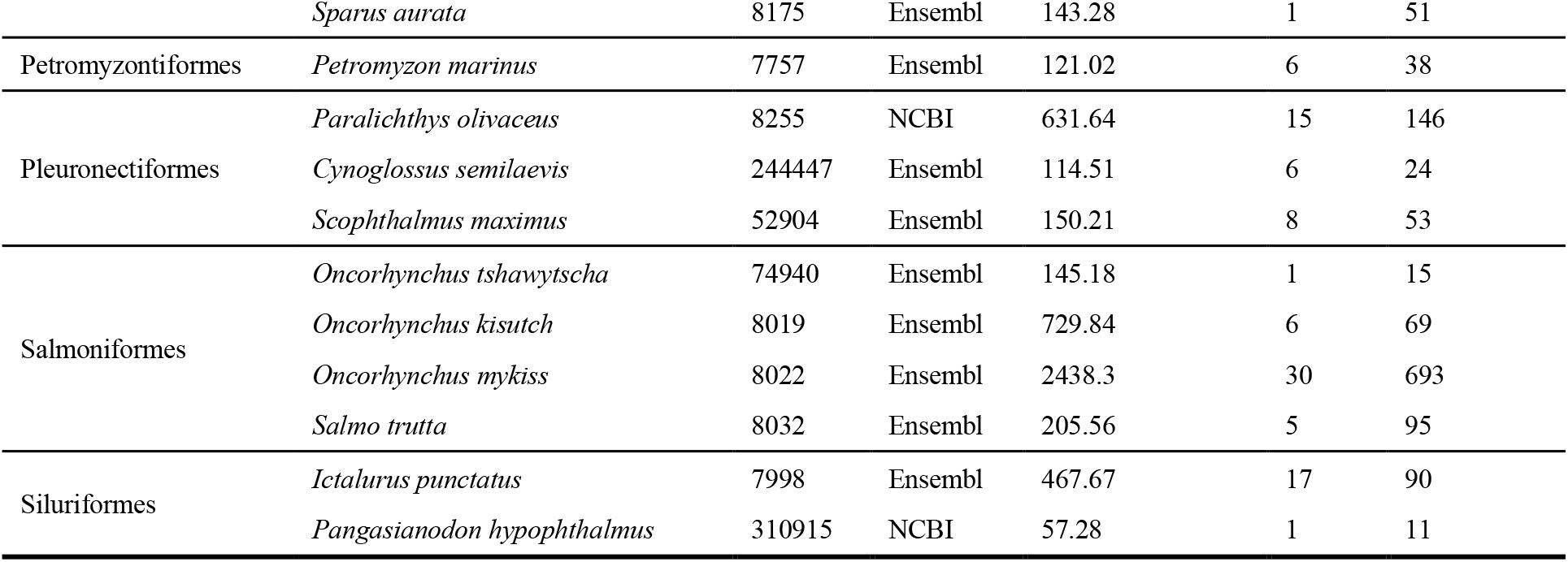
Fish species and corresponding data collected in FishExp.

The metadata is manually curated and organized, including strain, genotype, tissue, development and treatment. Figure 1 briefly illustrates the data collection, manual curation and data processing, and highlighted features of FishExp.

**Figure 1.**
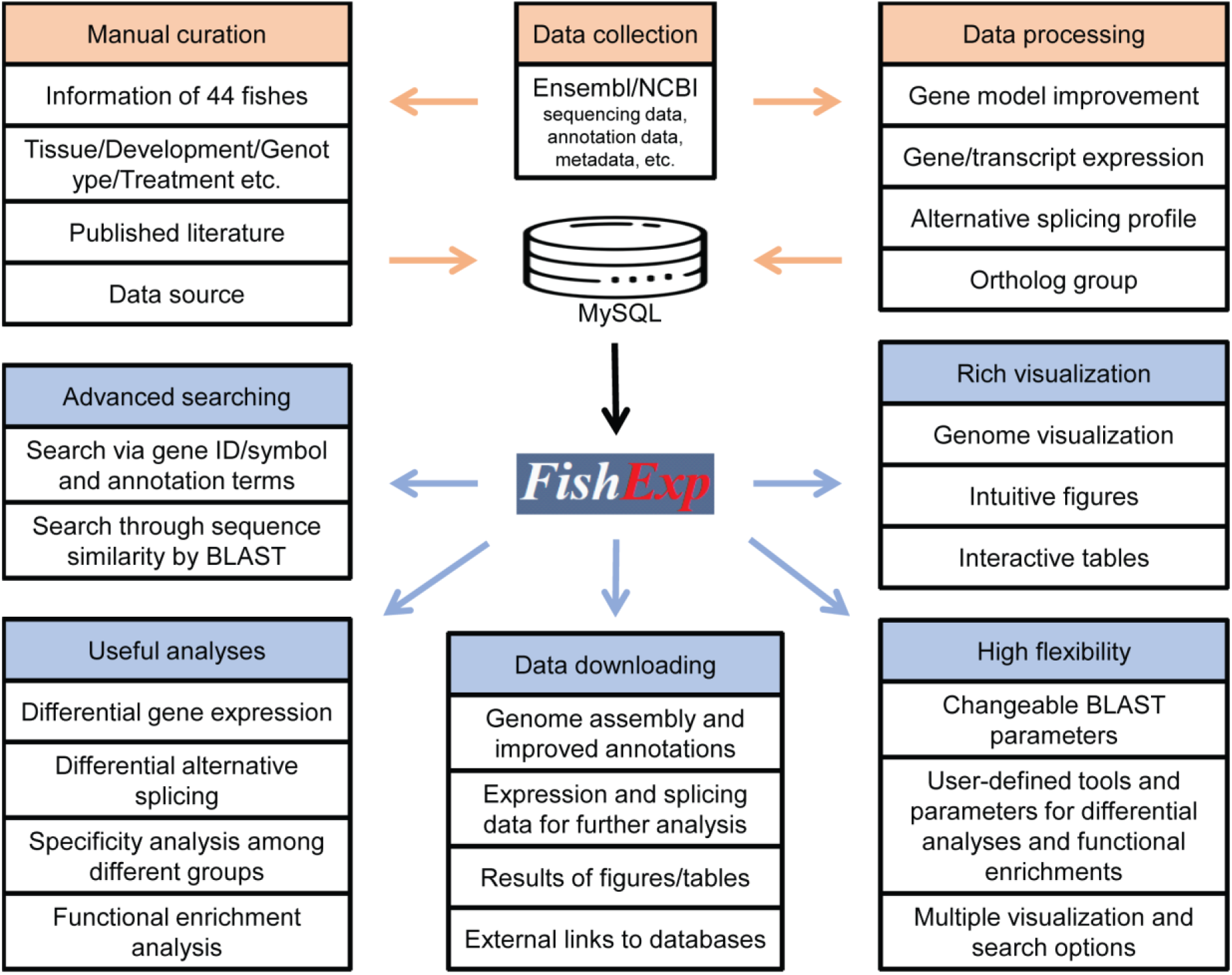
Overview of data collection, manual curation and data processing, and database features of FishExp.

### Gene annotation improvement

The reference genome sequences and annotation of 41 and 3 species are from the Ensembl and NCBI, respectively (Table 1). Accurate and complete gene annotation is extremely important for the unambiguous quantification of expression or splicing from RNA-seq experiments [18]. Whereas online genome annotations are largely incomplete, even for widely studied organism like human. In order to address this, we improved the gene/transcript model using the following steps. First, high-quality RNA-seq data sets were mapped to the reference genome by HISAT2 [19] and then assembled into transcripts using StringTie2 [20]. Second, we kept novel multi-exonic transcripts with at least 200 bps, an average coverage of 2x per transcript and 1x per exon. At last, we retained only the novel transcripts in at least 1/3 of all experiments and at least 3 experiments. All improved annotations are available for download on the Summary page of the FishExp database.

### Gene expression estimation and differential analysis

We used HISAT2 to perform read alignment against the reference genome and used StringTie2 to assembly them into transcripts and obtain expression levels for genes/transcripts. To perform differential expression analysis, two most widely used tools are employed: DESeq2 [21] and edgeR [22]. For edgeR, three tests are available including exact test, likelihood test and quasi-likelihood F test. It also supports comparison without sample replicates, which is useful for exploring many early sequencing experiments.

### Alternative splicing detection and differential analysis

By generating multiple isoforms from a single gene, AS influences diverse cellular processes including stability, localization, binding and enzymatic properties [23]. AS and DAS analyses could provide new insights into biological processes and disease conditions, however, they were underestimated and far from being fully mined in existing RNA-seq data sets. To overcome this, rMATs and customized scripts were used to perform analyses of AS and DAS. Five canonical AS types were considered, including exclusion or inclusion of individual exon (SE), alternative 5’ splice sites and 3’ splice sites (A5SS and A3SS), retention of intron (RI), mutually exclusive splicing of adjacent exons (MXE). Among them, SE is the most common type accounting for approximately 38.8% of all AS events detected in this study, followed by RI, A3SS and A5SS, while MXE only occurs in about 2.5% of AS events (Figure 2A). Approximately 13.4% of the total AS events were novel detected by rMATS based on read mapping (Figure 2B).

**Figure 2.**
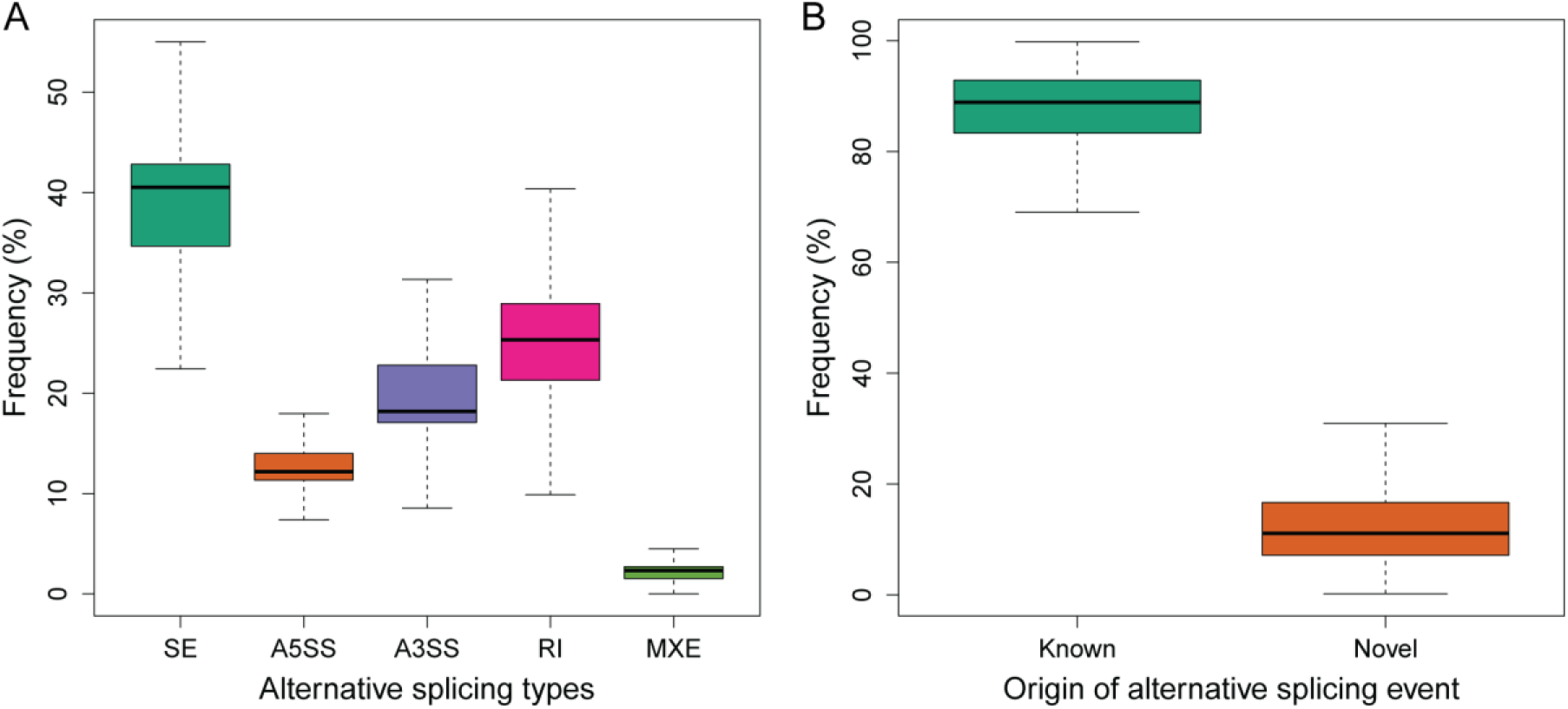
Summary of identified alternative splicing events. (**A**) Percentage of five common types of alternative splicing events in all collected RNA-seq experiments. SE, skipped exon; A5SS, alternative 5’ splice site; A3SS, alternative 3’ splice site; RI, retention of intron; MXE, mutually exclusive exon. (**B**) The detection of alternative splicing events is not only based on known transcripts, but also derived from high-quality read mapping against the corresponding genome.

### Functional enrichment

To help better understand the function of the biological system, we employed ClusterProfiler R package [24] to conduct functional enrichments of Gene Ontology (GO) and Kyoto Encyclopedia of Genes and Genomes (KEGG) pathway. For genomes from Ensembl, GO terms come from its functional annotation, while Blast2GO [25] was used to obtain GO annotations for RefSeq genomes. In addition, GO annotation of some species, especially model species, can also be directly extracted from AnnotationHub [26]. The pathway assignment was achieved by submitting protein sequence to KAAS (KEGG automatic annotation server) [27], which works mainly based on sequence similarities. The pathway annotation of widely studied species can be extracted directly from KEGG database. In addition to the typical enrichment analysis using hypergeometry test, we also provide options for enriching GO and pathways using Gene Set Enrichment Analysis (GSEA) [28].

### Orthologous gene and AS event

Gene conservation indicated positive natural selection, reflecting their evolutionary and functionally important roles. In an attempt to explore the gene conservation between fish species, orthologous genes between species were identified by OrthoFinder [29] using the longest protein sequence of each gene. Gene splicing can be found in almost all eukaryotic species. The formation and disappearance of AS events occur during evolution, and conserved AS provides strong evidence of biological function [30]. To explore the conservation of AS, we carried out the following steps. First, the protein sequences within one orthologous gene group were aligned using MAFFT [31]. Second, the protein alignments were converted to codon alignments using PAL2NAL [32]. Finally, we assigned new coordinates of exons in transcripts based on the codon alignments. All AS events with the same new coordinates were classified into an orthologous AS group. All the Orthologous genes and AS events were deposited in FishExp and had links to each other within one orthologous group.

### Implementation of FishExp

The data in FishExp are stored and managed in the relational databases MySQL. The web interfaces are based on HTML, CSS and JavaScript. The backend processing scripts use PHP, Perl and R language. A genome browser is implemented using JBrowser to facilitate convenient visualization of genes, transcripts and AS.

## Results

### Summary of genomic and transcriptomic data

The species table on the homepage lists all 44 species. The summary icon in the species table leads the user to view a brief biological introduction of the species, genomic statistics and the information of RNA-seq experiments. Of them, the genomic statistics include the number of original gene/transcript, improved transcript/exon/splice and AS event. Statistics of improvement results for all species are listed in Supplementary Table S1. On average, the number of exons and splice junctions increased by approximately 5.8% and 9.8%, respectively; the number of transcripts per multi-exon gene increased from 1.9 to 2.3; and the proportion of genes with alternative transcripts increased from 36.4% to 51.7% among all multi-exon genes. All the genomic and transcriptomic data, such as genome sequence/annotation, gene/transcript expression and AS data, are downloadable for users to conduct off-line analyses.

### Advanced searching

The search term for FishExp can be in any of the following formats: gene ID/symbol or functional categories including protein family, gene ontology and KEGG pathway (Figure 3A). In addition, the BLAST tools, including blastn, blastp and blastx, enable the user to search for targeted genes by supplying protein or nucleotide sequences from the current or other species (Figure 3B). The search results provide annotations with links to external mainstream databases including Swissport, pfam, GO and KEGG (Figure 3C). Furthermore, clicking on the expression tab will direct the user to the gene page, which displays 1) the general information of the gene, including the orthogroup with a popup window displaying the orthologous genes of other fish species, helping to explore the cross-species gene conservation; 2) a genome browser for intuitive visualization of the gene model and associated transcripts/AS event, from which the sequence of the region of interest can be obtained; 3) a hierarchical bar chart displaying gene expression profiles in experimental groups of a selected study at the first level and sequencing experiments at the second level by clicking a certain group. The gene page can lead the user to the transcript page and splicing page which have the similar layout (Figure 3D-G). Both transcript page and splicing page have links to each other and allow users to explore the potential functional impact of AS on associated transcripts (Figure 3H). Note that in the hierarchical bar chart of a selected study, the gene and transcript page present expression value using TPM (transcript per million) or FPKM (Fragments Per Kilobase of transcript per Million mapped reads) based on user selection, whereas the splicing page shows the percent spliced-in (PSI) indicating the exon inclusion level for a certain AS event.

**Figure 3.**
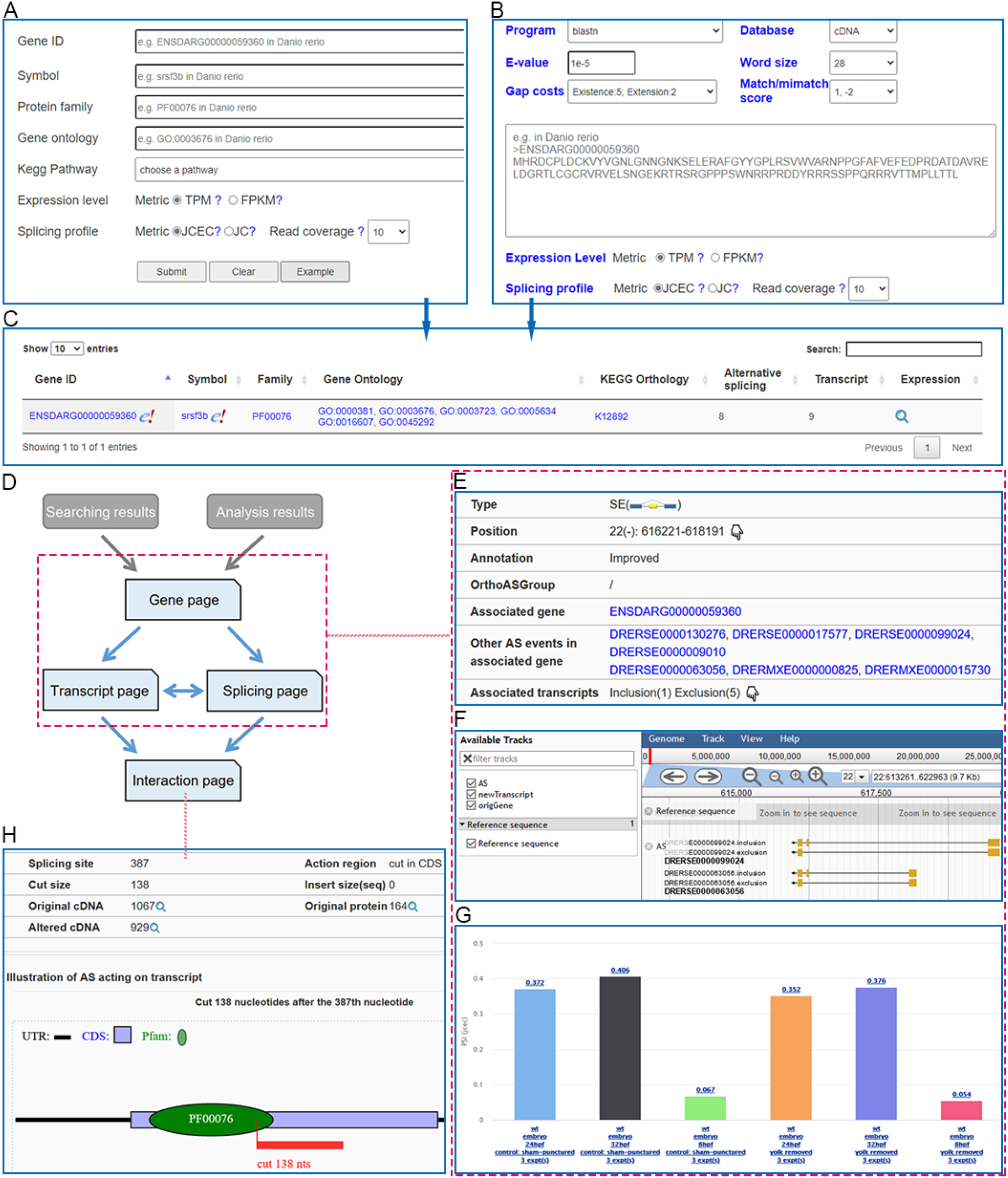
The advanced searching and display structure of FishExp. (**A**) Search: various options and filters for searching a gene of interest in a selected species. (**B**) BLAST: searching genes with user-inputted protein or nucleotide sequence from the current species or other species. Various parameters such as E-value and gap costs can be defined. (**C**) Searching results shows annotation information of the gene with external links and expression icon directing users to the gene page. (**D**) The display structure of FishExp. The gene page, transcript page and splicing page are interconnected and share similar webpage layout, including basic information, genome visualization and expression/splicing profiles, shown in (**E**) (**F**) (**G**) respectively, using splicing page as an example. (**H**) The interaction page illustrating the effect of an AS event on a selected transcript.

### Differential expression and alternative splicing

Most transcriptomic studies strive to identify and investigate differences between groups, which can provide insights into the underlying molecular mechanisms and generate new hypothesis. Here in the FishExp server, both differential gene expression (DGE) and differential alternative splicing (DAS) analyses are offered for users. The comparative analyses can be customized in many aspects. First, based on various information (strain, genotype, tissue, development and treatment), two groups of RNA experiments can be flexibly selected by clicking the corresponding icon in the “Control” and “Treatment” column (Figure 4A). In addition, the search box in the upper right corner can be used to locate certain study, RNA experiment or data source. Second, a number of analysis tools and parameters of DGE/DAS and GO/KEGG enrichment analyses are user-defined (Figure 4B and C). For instance, the DAS analysis provides different statistical models including MATS LRT, rMATS unpaired and rMATS paired.

**Figure 4.**
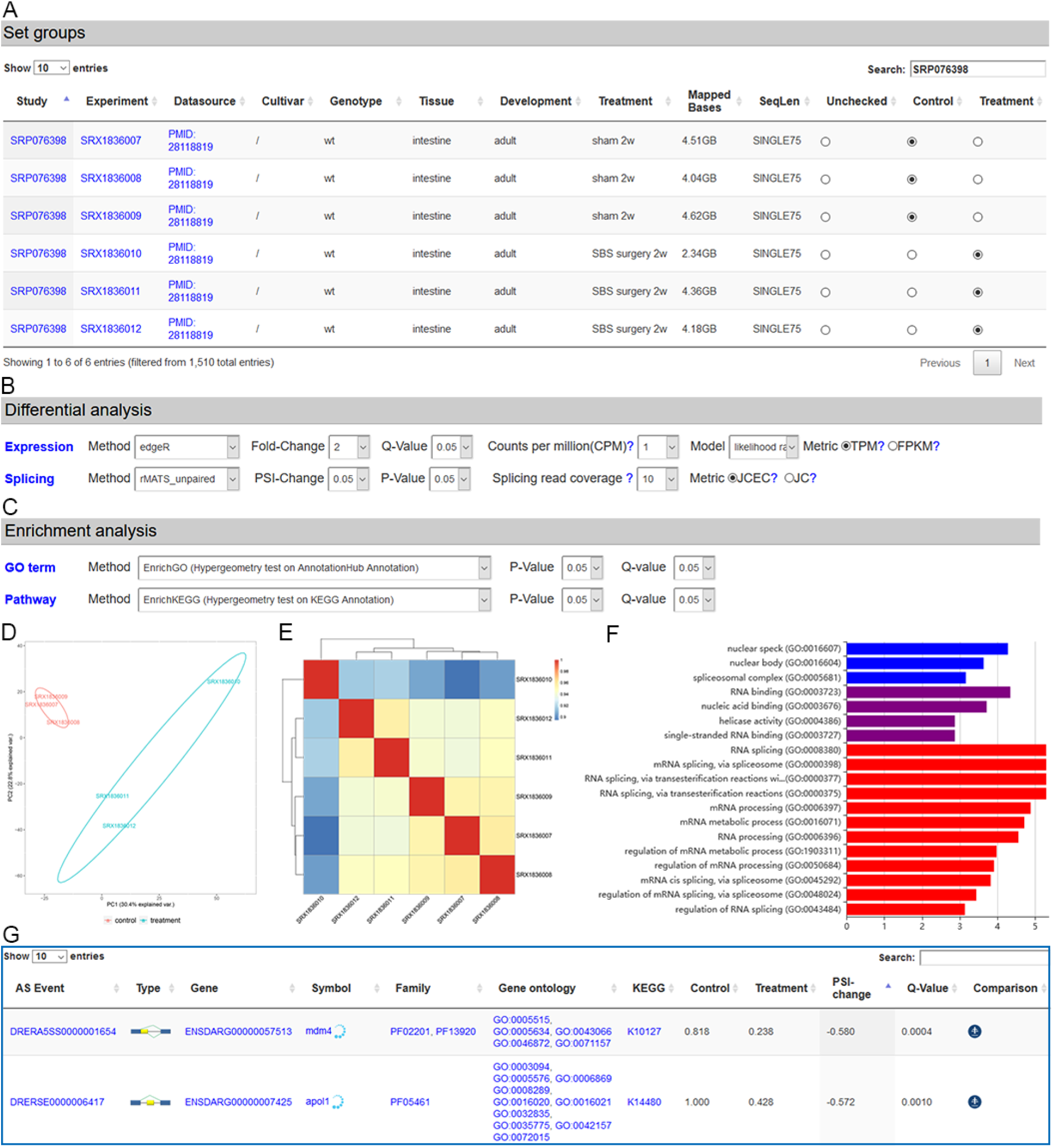
Comparative analyses in FishExp including differential expression and differential splicing. (**A**) Two groups, referred to as control and treatment, should be firstly selected by clicking on the corresponding icon. (**B**) Various tools, parameters and thresholds can be defined for the differential gene expression and alternative splicing analysis. (**C**) Functional enrichment can be conducted with many approaches and thresholds. (**D**) PCA plot and (**E**) heatmap showing the distribution of samples based on alternative splicing profiles. (**F**) Bar chart displaying the enrichment of GO terms. (**G**) The result table contains detailed information including links to other pages of FishExp and external databases.

The link of the result page including various tables and figures will be sent to the user-provided email after job submission. The top of the result page displays the basic analysis information of the user-selected samples, analysis tools and parameters. The remaining page is separated into two sections of analysis results: differential expression and differential splicing. Each section contains two clustering plots, PCA and heatmap (Figure 4D and E), which provide an overview of the variation of RNA-seq data and help check the group selection. It is followed by figures of enriched GO (Figure 4F) and pathway and the interactive table of DEG or DAS (Figure 4G) if available. Moreover, a download button is offered for users to obtain all the tables/high-quality figures for publication and other detailed information for further analysis.

### Specificity analysis

Specifically expressed genes (SEG) or specifically spliced genes (SSG) are of great interested since they may exert specific functions in a certain tissue, genotype or disease condition. The FishExp server provides the specificity analysis as an extended function of the differential analysis. The only difference is that at least four groups need to be selected in the specificity analysis, and the pairwise comparison between groups will be performed. The specifically expressed gene is referred to the gene which expressed significantly higher or lower in one group than that in any other groups. Similarly, the specifically alternatively spliced gene refers to the gene whose PSI value in one group is significantly higher or lower than that in other groups. The specificity result page is similar to that of the differential analysis between two groups, except that it additionally contains the chart displaying the number of SEG/SSG with significantly higher or lower expression/splicing levels than other groups.

### A case study for differential splicing patterns

We demonstrate here how the FishExp can reveal novel regulatory information from published research, promote the understanding of underlying mechanisms and help generate new hypothesis for future studies. A simple design in zebrafish was selected [33], which compared the gene expression in proximal intestine of adult zebrafish with and without short bowel syndrome (SBS). The SBS zebrafish model underwent treatment of laparotomy, proximal stoma and distal ligation, while the control fish experienced laparotomy alone. This study focused on overall changes in gene expression. We further performed the analyses of AS and DAS using FishExp web server with a few clicks and obtained the results in minutes (Figure 4).

Both PCA of gene expression and PCA of AS (Figure 4D) show that samples are distinguishably clustered to two groups in accordance to the treatment and control group, reflecting that SBS zebrafish exhibit evident changes in both gene expression and AS profiles. Using the selected tools and parameters (Figure 4B and C), a total of 76 DAS events of 66 DAS genes were identified, seven of which were also differentially expressed genes (Supplementary Table S2). Remarkably, the DAS genes were almost entirely enriched in the biological process of RNA splicing/processing and molecular function of RNA binding (Figure 4F). In addition, there is only one enriched KEGG pathway, spliceosome (dre03040), which plays central roles in the splicing process of eukaryotic genes. This observation is consistent with the findings that genes encoding RNA binding proteins including splicing factors and spliceosomal components themselves often undergo alternative splicing [34–39].

Splice site selection on pre-mRNA under specific conditions is determined by the binding of splicing factors, which recruit numerous spliceosomal components and thereby the spliceosome [40,41]. Changes in levels or activity of splicing factors may have profound effects on the expression of downstream target genes [37]. The main classes of splicing factors are Ser/Arg-rich (SR) proteins and heterogeneous nuclear ribonucleoprotein particle (hnRNP) proteins. Among the DAS events in this study, we identified four DAS events of three SR genes including *srsf3a* (SE and A3SS), *srsf4* (IR) and *srsf7a* (SE) as well as one differential A5SS event of *hnrnpr*. The expression or activity of these factors were regulated by alternative splicing, which may affect downstream gene network. For instance, the A5SS event on *hnrnpr* could result in an insertion of 18 nucleotides on the transcript of ENSDART00000172319, which overlap the translated acidic sequence segment domain (PF18360). The DAS events in the present study imply the involvement of alternative splicing in SBS, and the identified master regulators deserve further exploration.

## Discussion and future directions

The advent of RNA-seq has revolutionized our understanding in the complexity and function of gene expression regulation, and emphasized the considerable roles of AS in various biological processes. The study of AS is still lacking in fish species, whose RNA-seq data is sitting in public repertoire but not being fully explored. The hidden information may reveal valuable information on gene expression regulation and suggest new functional association for further investigations. We thereby created FishExp to help researchers address the complexity in analyzing and visualizing the gene expression and alternative splicing profiles.

Serving as an added-value resource, FishExp not only provides a comprehensive survey of the profiles of gene expression/splicing, but also is ideally suited to study the DEG and DAS genes/events and investigate their functional roles: 1) The comparison analysis allows user to study differential genes between two groups, and 2) the specificity analysis is designed to explore specifically expressed or spliced genes in each group relative to others. These differential and specific genes/events may indicate crucial biological functions, as such, the offered analyses would be of great interest to a broad range of users. In addition to provide a wealth of information and analysis tools, we sought to make the database easy to use. The database is easy to navigate with logical, hierarchical and interactive webpage, displayed with a highly interactive interface. Many external links are embedded for further exploration, and all data can be downloaded in batch for additional offline analysis.

In the future, we strive to frequently update FishExp to cover newly generated transcriptome data for current and additional fish species, including long sequencing reads for accurate transcript assembly and splicing detection. We envision to extend FishExp to provide more analyses, such as alternative polyadenylation and RNA-editing. We anticipate FishExp to become a very useful resource to explore the profiles and functions of gene expression/splicing, to expand our understanding of gene expression regulation, and to promote hypothesis generation for further research.

### Key points

We developed FishExp, which enables retrieval and integrated analysis of gene expression and alternative splicing profiles, processed from 26,081 RNA-seq experiments of 664 studies of 44 fish species, using a uniform pipeline.

All samples were manually curated to label their information, including strains, genotypes, tissues, developmental stages and experimental conditions. Additionally, orthologous gene and alternative splicing groups were identified and presented.

Differential/specific gene expression and differential/specific alternative splicing analyses are offered using customized experiments and tools, and functional enrichments are available to assign biological significance.

FishExp provides a wide range of visualization tools and a user-friendly interface with gene-, transcript- and splicing event-level webpage. Additionally, all data can be downloaded for further analysis.

## References

1. Froese R, Pauly D. FishBase in the Catalogue of Life. 2019.

2. Friedman M, Sallan LC. Five hundred million years of extinction and recovery: a Phanerozoic survey of large‐scale diversity patterns in fishes. Palaeontology 2012;55:707–742.

3. Shu D-G, Morris SC, Han J, et al. Head and backbone of the Early Cambrian vertebrate Haikouichthys. Nature 2003;421:526–529.

4. Meyers JR. Zebrafish: development of a vertebrate model organism. Curr. Protoc. Essent. Lab. Tech. 2018;16:e19.

5. Ablain J, Zon LI. Of fish and men: using zebrafish to fight human diseases. Trends Cell Biol. 2013;23:584–586.

6. Schartl M. Beyond the zebrafish: diverse fish species for modeling human disease. Dis. Model. Mech. 2014;7:181–192.

7. Béné C, Arthur R, Norbury H, et al. Contribution of fisheries and aquaculture to food security and poverty reduction: assessing the current evidence. World Dev. 2016;79:177–196.

8. Qian X, Ba Y, Zhuang Q, et al. RNA-Seq Technology and Its Application in Fish Transcriptomics. Omi. A J. Integr. Biol. 2014;18:98–110.

9. Pan Q, Shai O, Lee LJ, et al. Deep surveying of alternative splicing complexity in the human transcriptome by high-throughput sequencing. Nat. Genet. 2008;40:1413–1415.

10. Wang ET, Sandberg R, Luo S, et al. Alternative isoform regulation in human tissue transcriptomes. Nature 2008;456:470–476.

11. Ule J, Blencowe BJ. Alternative splicing regulatory networks: functions, mechanisms, and evolution. Mol. Cell 2019;76:329–345.

12. Ryan M, Wong WC, Brown R, et al. TCGASpliceSeq a compendium of alternative mRNA splicing in cancer. Nucleic Acids Res. 2016;44:D1018–D1022.

13. Hyung D, Kim J, Cho SY, et al. ASpedia: a comprehensive encyclopedia of human alternative splicing. Nucleic Acids Res. 2018;46:D58–D63.

14. Tapial J, Ha KCH, Sterne-Weiler T, et al. An atlas of alternative splicing profiles and functional associations reveals new regulatory programs and genes that simultaneously express multiple major isoforms. Genome Res. 2017;27:1759–1768.

15. Liu J, Tan S, Huang S, et al. ASlive: a database for alternative splicing atlas in livestock animals. BMC Genomics 2020;21:1–7.

16. Li Z, Zhang Y, Bush SJ, et al. MeDAS: a Metazoan Developmental Alternative Splicing database. Nucleic Acids Res. 2021;49:D144–D150.

17. Zhu Y, Stephens RM, Meltzer PS, et al. SRAdb: query and use public next-generation sequencing data from within R. BMC Bioinformatics 2013;14:1–4.

18. Zhang D, Guelfi S, Garcia-Ruiz S, et al. Incomplete annotation has a disproportionate impact on our understanding of Mendelian and complex neurogenetic disorders. Sci. Adv. 2020;6:eaay8299.

19. Kim D, Paggi JM, Park C, et al. Graph-based genome alignment and genotyping with HISAT2 and HISAT-genotype. Nat. Biotechnol. 2019;37:907–915.

20. Kovaka S, Zimin A V, Pertea GM, et al. Transcriptome assembly from long-read RNA-seq alignments with StringTie2. Genome Biol. 2019;20:1–13.

21. Love MI, Huber W, Anders S. Moderated estimation of fold change and dispersion for RNA-seq data with DESeq2. Genome Biol. 2014;15:550.

22. Robinson MD, McCarthy DJ, Smyth GK. edgeR: a Bioconductor package for differential expression analysis of digital gene expression data. Bioinformatics 2010;26:139–140.

23. Kelemen O, Convertini P, Zhang Z, et al. Function of alternative splicing. Gene 2013;514:1–30.

24. Yu G, Wang LG, Han Y, et al. ClusterProfiler: An R package for comparing biological themes among gene clusters. Omi. A J. Integr. Biol. 2012;16:284–287.

25. Conesa A, Götz S. Blast2GO: a comprehensive suite for functional analysis in plant genomics. Int. J. Plant Genomics 2008;2008:619832.

26. Morgan M, Carlson M, Tenenbaum D, et al. Package ‘AnnotationHub’. 2019.

27. Moriya Y, Itoh M, Okuda S, et al. KAAS: an automatic genome annotation and pathway reconstruction server. Nucleic Acids Res. 2007;35:W182–W185.

28. Subramanian A, Tamayo P, Mootha VK, et al. Gene set enrichment analysis: a knowledge-based approach for interpreting genome-wide expression profiles. Proc. Natl. Acad. Sci. 2005;102:15545–15550.

29. Emms DM, Kelly S. OrthoFinder: phylogenetic orthology inference for comparative genomics. Genome Biol. 2019;20:1–14.

30. Keren H, Lev-Maor G, Ast G. Alternative splicing and evolution: diversification, exon definition and function. Nat. Rev. Genet. 2010;11:345.

31. Katoh K, Standley DM. MAFFT multiple sequence alignment software version 7: improvements in performance and usability. Mol. Biol. Evol. 2013;30:772–780.

32. Suyama M, Torrents D, Bork P. PAL2NAL: robust conversion of protein sequence alignments into the corresponding codon alignments. Nucleic Acids Res. 2006;34:W609–W612.

33. Schall KA, Thornton ME, Isani M, et al. Short bowel syndrome results in increased gene expression associated with proliferation, inflammation, bile acid synthesis and immune system activation: RNA sequencing a zebrafish SBS model. BMC Genomics 2017;18:1–13.

34. Saltzman AL, Pan Q, Blencowe BJ. Regulation of alternative splicing by the core spliceosomal machinery. Genes Dev. 2011;25:373–384.

35. Tan S, Wang W, Zhong X, et al. Increased Alternative Splicing as a Host Response to Edwardsiella ictaluri Infection in Catfish. Mar. Biotechnol. 2018;1–10.

36. Lareau LF, Inada M, Green RE, et al. Unproductive splicing of SR genes associated with highly conserved and ultraconserved DNA elements. Nature 2007;446:926–929.

37. Staiger D, Brown JWS. Alternative splicing at the intersection of biological timing, development, and stress responses. Plant Cell 2013;25:3640–3656.

38. Zhang Y, Wu X, Li J, et al. Comprehensive characterization of alternative splicing in renal cell carcinoma. Brief. Bioinform. 2021.

39. Phillips JW, Pan Y, Tsai BL, et al. Pathway-guided analysis identifies Myc-dependent alternative pre-mRNA splicing in aggressive prostate cancers. Proc. Natl. Acad. Sci. 2020;117:5269–5279.

40. Matlin AJ, Clark F, Smith CWJ. Understanding alternative splicing: towards a cellular code. Nat. Rev. Mol. cell Biol. 2005;6:386–398.

41. Nilsen TW, Graveley BR. Expansion of the eukaryotic proteome by alternative splicing. Nature 2010;463:457–463.

